# Determining sufficient sequencing depth in RNA-Seq differential expression studies

**DOI:** 10.1101/635623

**Authors:** Andrew J. Bass, David G. Robinson, John D. Storey

## Abstract

RNA-Seq studies require a sufficient read depth to detect biologically important genes. Sequencing below this threshold will reduce statistical power while sequencing above will provide only marginal improvements in power and incur unnecessary sequencing costs. Although existing methodologies can help assess whether there is sufficient read depth, they are unable to guide how many additional reads should be sequenced to reach this threshold. We provide a new method called superSeq that models the relationship between statistical power and read depth. We apply the superSeq framework to 393 RNA-Seq experiments (1,021 total contrasts) in the Expression Atlas and find the model accurately predicts the increase in statistical power gained by increasing the read depth. Based on our analysis, we find that most published studies (> 70%) are undersequenced, i.e., their statistical power can be improved by increasing the sequencing read depth. In addition, the extent of saturation is highly dependent on statistical methodology: only 9.5%, 29.5%, and 26.6% of contrasts are saturated when using `DESeq2`, `edgeR`, and `limma`, respectively. Finally, we also find that there is no clear minimum per-transcript read depth to guarantee saturation for an entire technology. Therefore, our framework not only delineates key differences among methods and their impact on determining saturation, but will also be needed even as technology improves and the read depth of experiments increases. Researchers can thus use superSeq to calculate the read depth to achieve required statistical power while avoiding unnecessary sequencing costs.

## Introduction

RNA-Seq technology is often used to measure genome-wide gene expression from multiple biological conditions to identify differentially expressed genes. There are two primary sources of variation in an experiment that impact the ability to detect differentially expressed genes: the biological variation of gene expression inherent in cellular samples and the technical variation induced once the samples are collected, processed, and sequenced. While biological variation can be reduced by increasing the number of biological samples included in the study, technical variation can be reduced through improved processing protocols and the manner in which the resulting libraries are sequenced. In particular, technical variation can be controlled through the number of reads sequenced (i.e., read depth). Given that current sequencing technologies provide a broad range of possible read depths, choosing the read depth in a study requires careful consideration with respect to statistical power, sample size, and cost [1, 2]. Sequencing too few reads will lead to unreasonably high technical variation and reduced statistical power, whereas sequencing too many reads will provide only marginal improvements in statistical power and incur unnecessary costs. Therefore, an important objective when utilizing sequencing technology is to determine a sufficient read depth that achieves acceptable statistical power without incurring unnecessary experimental costs [3, 4]. However, there are currently no methods to guide researchers in determining how many additional reads should be obtained in an RNA-Seq study.

There are several heuristic methodologies for assessing the relationship between sequencing depth and statistical power within the sequencing depth range of a completed experiment [3, 5, 6]. These approaches are commonly referred to as ‘subsampling’ (or ‘downsampling’) procedures, where experiments are simulated at lower read depths by randomly sampling reads from the completed experiment. For each simulated experiment, the number of differentially expressed genes at a specified false discovery rate (FDR) is calculated to determine the effect of lower read depths on statistical power. When applying these methods, the goal is to assess whether or not the experiment is near a saturation point, where increasing the read depth will only provide marginal improvements in statistical power. As an illustration, we applied the subsampling methodology `subSeq` [6] to two experiments involving *Saccharomyces cerevisiae* that have different sequencing depths (Figure 1a-b). In Figure 1a, the experiment is oversaturated and sequencing additional reads will provide only marginal improvements in power while incurring unnecessary costs. Alternatively, in Figure 1b, the experiment is undersaturated and sequencing additional reads will improve the statistical power. While subsampling methods are informative, they are unable to estimate the number of additional reads needed to reach saturation for undersaturated experiments.

We provide a solution to this problem by modeling and predicting the relationship between sequencing depth and statistical power in a completed experiment. Our method superSeq can accurately predict how many additional reads, if any, need to be sequenced in order to maximize statistical power given the number of biological samples. Applied to the above example, our model (dashed line) predicts, as expected, that sequencing more reads in Figure 1a (oversaturation) provides marginal improvements in the number of differentially expressed genes detected. Conversely, in Figure 1b (undersaturation), substantially more differentially expressed genes will be found by doubling the read depth. Therefore, the predictions from our model not only indicate when an experiment has sufficient read depth, but can also be used to inform how many additional reads should be sequenced when an experiment is undersaturated. An important point to note is that the saturated experiment has substantially lower read depth and the same number of biological replicates compared to the undersaturated experiment, even though both involve the same organism. This suggests that determining sufficient read depth is not simply organism-specific but may vary for every experimental design.

**Figure 1:**
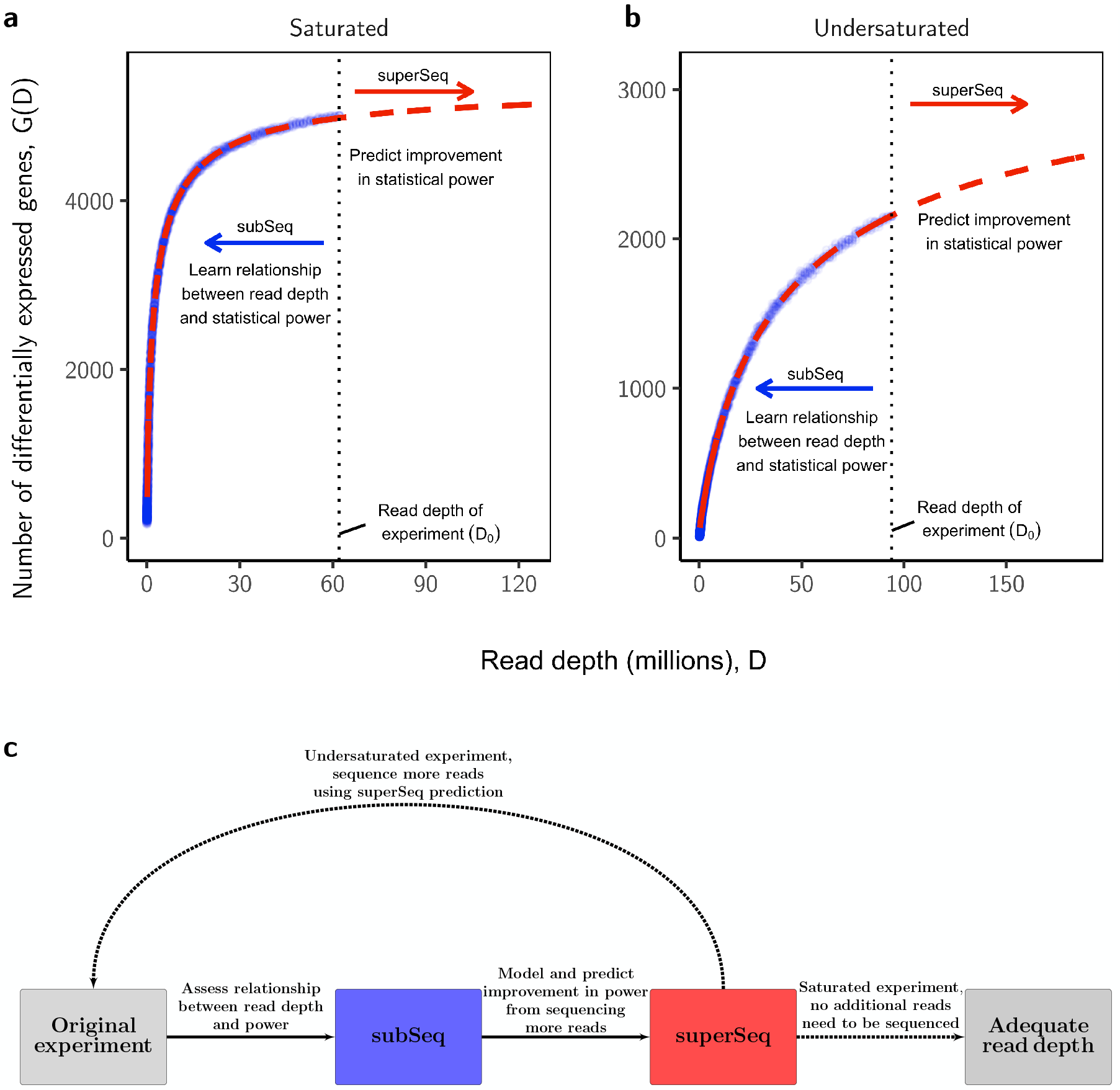
Learning the relationship between statistical power and sequencing depth. **(a,b)** The superSeq model was applied to two different experiments that were either saturated or undersaturated with reads, and the differential expression method `DESeq2` was used to calculate the number of differentially expressed genes at various subsampling proportions as shown by the points. The vertical dotted lines indicate the sequencing depth of the experiment and the dashed lines are the predictions from superSeq. **(c)** A process to determine sufficient read depth in an RNA-Seq experiment: assess the relationship between read depth and statistical power (blue), use superSeq to model and predict the improvement in power from sequencing additional reads (red). If the experiment is undersaturated then additional reads should be sequenced using the predictions from superSeq.

## Models and methods

The shapes of the superSeq curves in Figure 1 are determined by the test statistics, the read depth, the number of biological replicates, and a prespecified FDR. Once the experiment has been completed and the number of biological replicates is fixed, we model the distribution of test statistics as a function of the read depth *D*. In particular, the data from subsampling is utilized to develop insights into how the test statistics vary at lower read depths. Our model then leverages this information to predict statistical power at higher read depths. More specifically, consider a completed sequencing experiment with initial read depth *D*_0_. We can parametrically model the number of differentially expressed genes detected at read depth proportion 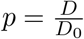 as

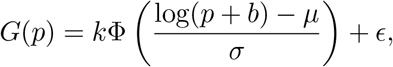

where Φ is the Normal cumulative distribution function with mean *μ* and variance *σ*^2^; *k* is the expected number of differentially expressed genes when the technical variation is minimized (with respect to a prespecified FDR); *b* is an offset parameter; and *ϵ* is a Normal random error (detailed in Appendix B). Thus *G*(*p* = 1) is the observed number of differentially expressed genes at the original sequencing depth *D*_0_. We obtain simulated observations of *G*(*p*) at lower sequencing depths (*p* < 1) by applying the subSeq methodology [6]. Using these observations, the above parameters in our model are estimated by implementing a non-linear least squares algorithm. The fitted curve is then extrapolated beyond the original read depth (*p* > 1) to provide predictions.

**Figure 2:**
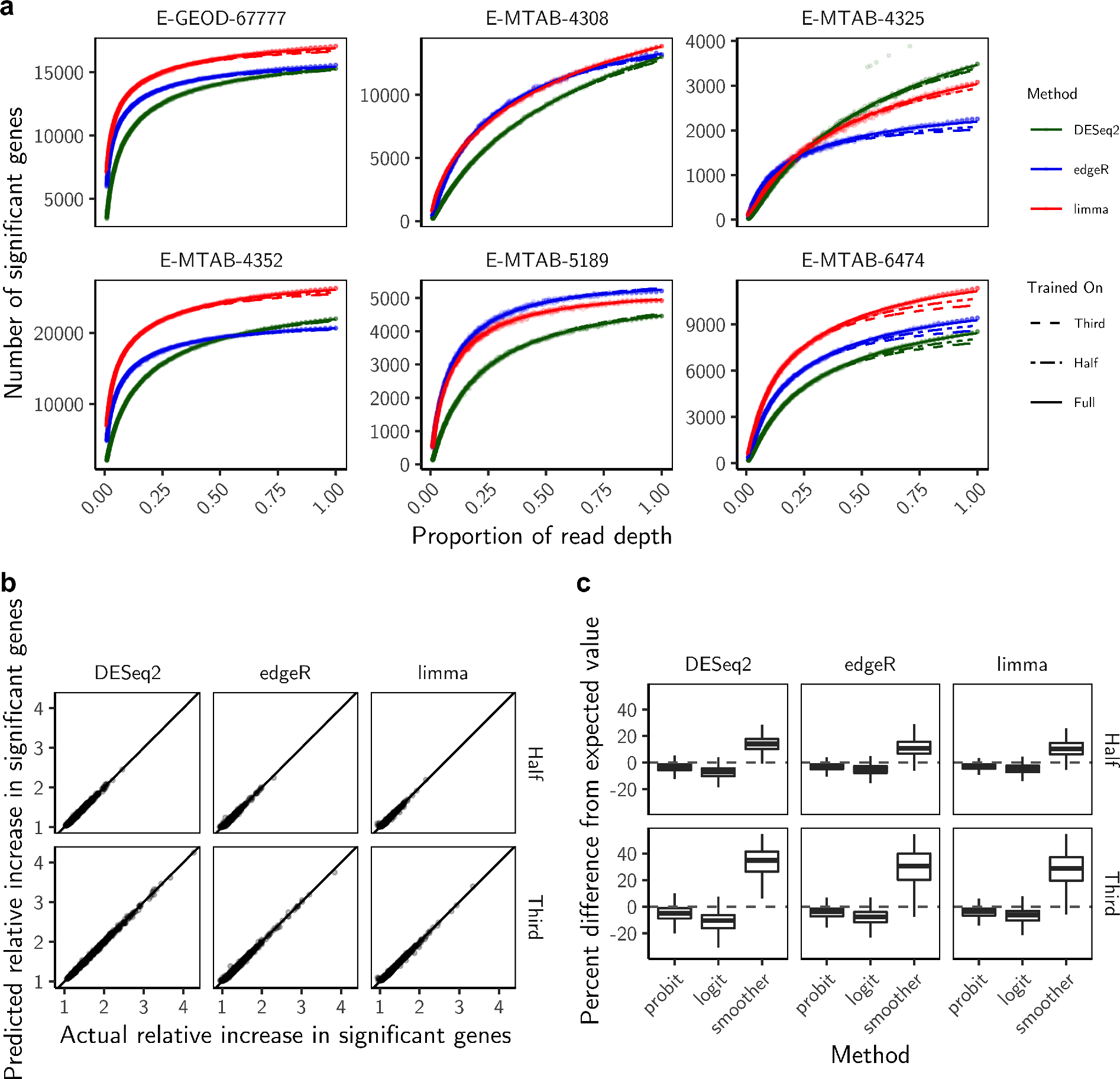
Validation of proposed model on real experimental data. **(a)** The number of significant genes at each depth, along with predicted parametric curves using superSeq for six randomly selected experimental contrasts. Each superSeq fit was performed three times: on the full experiment and on data that had previously been presampled to one-third and one-half of the full read depth. The differential expression packages `DESeq2`, `edgeR`, and `limma` were used to test for differential expression at each subsampling proportion. Using the estimates from these packages, we applied our superSeq model to predict the increase in power from an increase in read depth. **(b)** The predicted relative increase in significant genes at the full read depth from training superSeq on a half and a third of the reads compared to the actual relative increase in significant genes detected at the full read depth. Each point represents a different experimental contrast in the Expression Atlas. In total, there are 1,021 experimental contrasts plotted. The points along the line represent accurate estimates of the predicted values. **(c)** Comparison of three different model fits to predict the relationship between statistical power and sequencing depth: probit (used in superSeq), logit, and smoother. We applied each model to all experimental contrasts and tested the predictive power when the curve is extrapolated to the full read depth.

## Results

To validate the model, we performed the following study on 393 RNA-Seq experiments from the Expression Atlas [7]. For each experiment, we trained the superSeq model on one-third and one-half of the full read depth, and then extrapolated the resulting fit to the full read depth to compare our model predictions with the observed experiment (Figure 1c). In this way, we are utilizing the sequenced read depth and associated statistical power of each experiment to test the accuracy of our model. Although any differential expression method can be used within the superSeq framework, we applied a weighted least squares regression (`limma` [8]) and two different implementations of a Negative Binomial regression (`edgeR` [9] and `DESeq2` [10]) to all experiments (detailed in Appendix A). We find that our model accurately predicts the increase in statistical power with increasing read depth, and in many cases is almost indistinguishable from the fit using the full read depth (Figure 2a). Additionally, we find that our predictive model provides a useful metric for estimating saturation: across the majority of experiments, using either differential expression method, the model accurately predicts statistical power at the full read depth, even when trained on just one-third of the reads (Figure 2b).

We compared the performance of our model (labeled ‘probit’) to two alternative approaches, namely, a cubic smoothing spline (labeled ‘smoother’) and a logistic distribution with the same number of free parameters (labeled ‘logit’) in Figure 2c. Natural cubic splines are known to predict poorly outside the range of observed data; as expected, it overestimates the sufficient read depth. While the logistic distribution has a similar shape (S-shaped) to our model, we find that the logit model substantially underestimates the sufficient read depth. These observations suggest that our model is most accurately capturing the observed relationship between sequencing depth and statistical power across all experiments in these comparisons.

The RNA-Seq experiments show varying degrees of saturation of both significance and accuracy (Figure 3a). For example, some experiments were able to detect 75% of the differentially expressed genes from the full experiment even when read depth had been reduced to 10% of its total, while others detected less than 5% of the genes at this read depth. Similarly, Spearman correlations between effect size estimates at this depth and the actual estimates varied from 0.5 to 0.9. This indicates that some experiments were saturated with reads, and that their read depth could have been substantially reduced while still achieving similar statistical power. In turn, experiments with a nearly linear increase in statistical power up to the full read depth suggest that the read depth of the experiment was insufficient, and increasing the read depth further would lead to greater statistical power.

**Figure 3:**
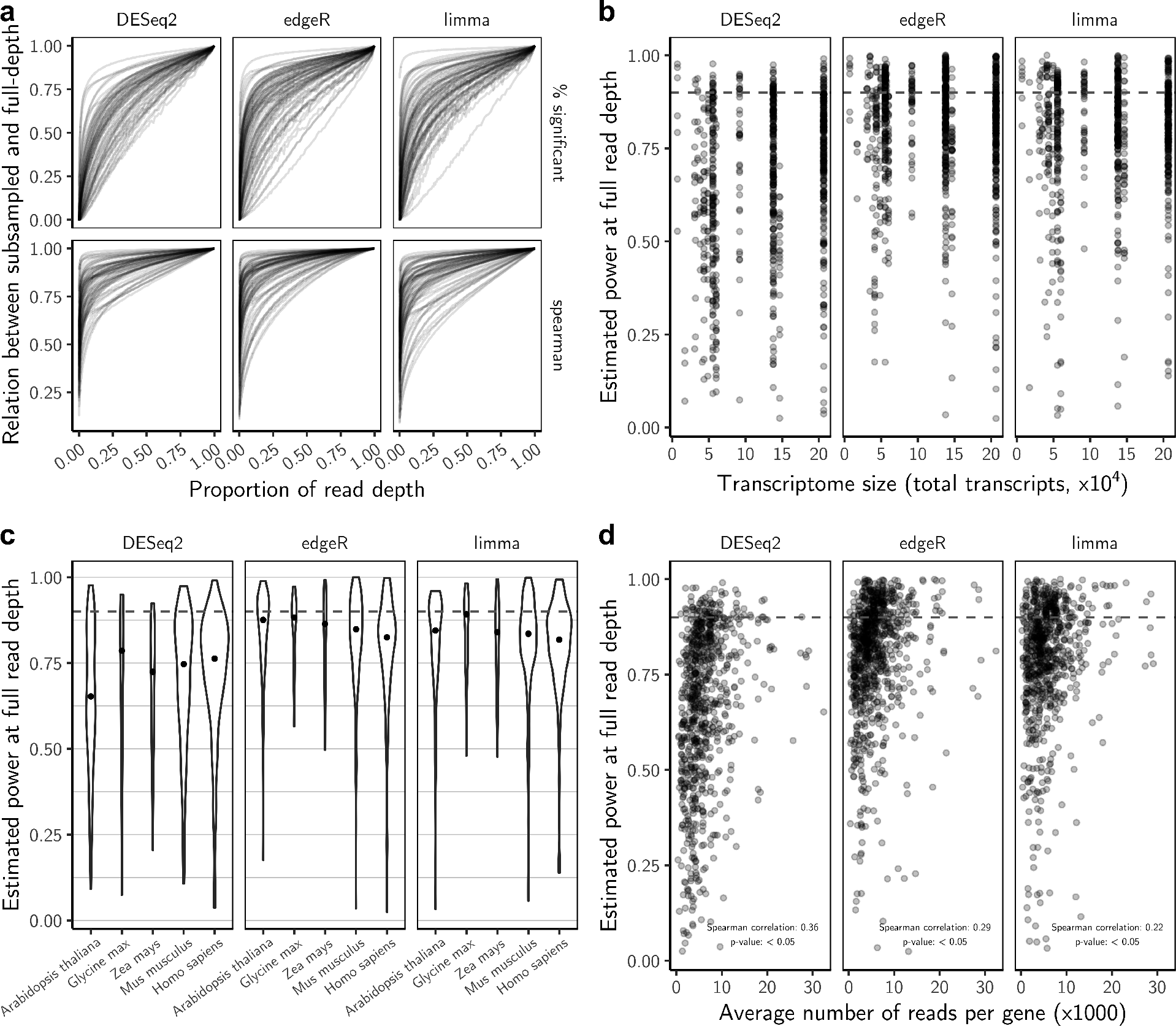
Quantifying the extent of saturation across multiple organisms. **(a)** Metrics showing power across varying read depths for 75 randomly chosen differential expression contrasts, using the software packages `DESeq2`, `edgeR`, and `limma` to test for differential expression. Each value was averaged across six replicates from subsampling. **(b)** Relationship between transcriptome size and the extent of saturation. For each experiment, we trained superSeq on the full read depth and the estimated power at the original sequencing depth was calculated using the model fit. Estimated power values above the dashed line at 0.9 (90% power) indicate that the experiment is near saturation while points below are undersaturated. **(c)** We selected five species that have the most experiments in the Expression Atlas to assess saturation between species. The black point is the median value. **(d)** Relationship between the average number of reads per gene and the extent of saturation (using the estimated power at the full read depth) across the Expression Atlas experiments. Each point represents a different experiment and values below the dashed line are undersaturated while points above are saturated.

We quantified the extent of saturation by calculating the estimated power from our model fit at the observed read depth for each experiment. We find the majority of experiments were undersequenced with reads and the extent of saturation varied depending on which differential expression method was applied. Specifically, only 9.5%, 29.5%, and 26.6% of experiments had sufficient read depth (defined in this instance as estimated power greater than 90% at the observed read depth) when using `DESeq2`, `edgeR`, and `limma`, respectively. We then tested whether the extent of saturation varied with respect to transcriptome size and find a statistically significant negative correlation when using `edgeR` (Spearman correlation: −0.07; *p* < 0.05) and a statistically significant positive correlation when using `DESeq2` (Spearman correlation: 0.18; *p* < 0.05). However, the variation in saturation at similar transcriptome sizes is comparatively large (Figure 3b). Furthermore, when grouping experiments by organism, we find that there is no clear trend in the extent of saturation across species. For example when using `limma` and `edgeR`, experiments performed on *Arabidopsis thaliana* were marginally more saturated than experiments performed on *Homo sapiens*, which has approximately four times the transcriptome size (Figure 3c). Conversely, when using `DESeq2`, the experiments performed on *Homo sapiens* were more saturated compared to those on *Arabidopsis thaliana*.

One important question is whether saturation in an experiment can be determined by a simpler approach, such as using the average number of reads per gene. We examined how this value is related to the extent of saturation (Figure 3d). As expected, there is a statistically significant negative correlation (*p* < 0.05) between the per gene read depth and the extent of saturation in each method. However, the variation in the degree of saturation between experiments at similar read depths is relatively large. Furthermore, experiments with large per gene read depths are less saturated than experiments at lower per gene read depths. These results suggest that one cannot assign a simple minimum per gene read depth threshold for an entire technology.

## Conclusions

Prior to this work, researchers designing experiments have had to rely on similar studies to guide their choice of read depth [1, 11, 12, 13]; however, finding such studies can often be challenging. Furthermore, methodologies available to assess the relationship between read depth and statistical power do not guide researchers in predicting a sufficient read depth. Our model, superSeq, can be used with any completed experiment to predict the relationship between statistical power and read depth. This will allow researchers to support their biological conclusions by demonstrating their experiment has adequate statistical power. In addition, our method can also be used as a diagnostic tool to model technical variation in a study when there are unexpected biological findings.

We find that superSeq provides accurate predictions when applied to real experiments with varying degrees of saturation, and that the extent of saturation in experiments depends on the differential expression method. This suggests that the estimated technical variability in an experiment varies with methodology: `DESeq2` generally estimates higher dispersion than `edgeR` or `limma`. Additionally, the majority of studies in Expression Atlas, regardless of the differential expression method, were undersaturated with reads, suggesting that these experiments could benefit from sequencing additional reads, as shown from the superSeq predictions. Finally, there was not a clear guideline when assigning a minimum per gene read depth to guarantee saturation for an entire technology, and thus superSeq will still be needed even as technology improves and the read depth of experiments increases.

Although only two-sample comparisons were used in this study, the model can be extended to any complex experimental design, such as time course studies [14]. We are currently extending superSeq to other areas where sufficient sequencing coverage is essential. In ChIP-Seq experiments, a common issue is determining the optimal coverage for differential binding analysis. In single-cell RNA-Seq experiments, adequate coverage is important when testing for differential expression across different populations of cells. This will also allow for a more comprehensive analysis of the model.

## Software and reproducible analysis

The data and code used to generate the results in this paper can be found at https://github.com/StoreyLab/superSeq-manuscript. An implementation of our method is available in the R package `superSeq` and can be found at https://github.com/StoreyLab/superSeq.

## Appendix

### A Application to Expression Atlas

The experiments used in the paper were from an online repository of gene expression experiments called Expression Atlas [7]. From the repository, we obtained 465 RNA-Seq differential expression experiments across 39 organisms. Each experiment had at least one two-sample comparison and there were a total of 1,353 comparisons. For each two-sample comparison, we first filtered out genes with fewer than 5 reads, then used the R package `subSeq` to subsample the datasets. Although any sub-sampling method can be applied, `subSeq` is a memory efficient and computationally fast implementation [6]. We subsampled these datasets across 301 subsampling proportions spaced on a logarithmic scale from 0.001 to 1. There were six replications at each subsampling proportion, and we used `edgeR`, `DESeq2`, and `limma` to test for differential expression [8, 9, 10]. Significant genes were determined at a *q*-value cutoff of 0.05 using the `qvalue` package [15].

We filtered out subsampling curves where fewer than 100 differentially expressed genes were detected at one-third of the read depth: the signal-to-noise ratio of these filtered experiments were too low to test the model. After filtering, there were 1,045, 998, and 919 subsampling curves for `edgeR`, `DESeq2`, and `limma`. We also removed subsampling curves that were non-monotonic and/or highly unstable. After this filtering step, there were 981, 859, and 851 subsampling curves for `edgeR`, `DESeq2`, and `limma`. In total, there were 393 experiments and 1,021 two-sample comparisons used to train our model. We then created two additional datasets for each study to test our model: one at one-third and the other at one-half of the full read depth using the observations from `subSeq`.

We fit our proposed model to each dataset and extrapolated the fit to the full read depth of the original experiment. There were a few studies with unstable subsampling curves at low subsampling proportions. Therefore, our model was trained on subsampling proportions 0.01 to 1. We then used the R package `nls` to fit a nonlinear least squares algorithm with the following parameters: 10,000 iterations, a convergence criterion of 10^−6^, and a minimum step size of 0.0002.

As stated in the main text, the Expression Atlas data and code used to generate the figures in this paper can be found at https://github.com/StoreyLab/superSeq-manuscript. The experiment accession IDs used to produce Figure 1 were E-MTAB-5313 (labeled Saturated) and E-GEOD-59814 (labeled Undersaturated). There were 3 biological replicates for each condition and the observed read depths were 62,085,792 and 94,023,022, respectively.

### B Relationship between the number of rejections and read depth

Under a Bayesian mixture model, the positive false discovery rate (pFDR) at significance threshold λ for test statistic *τ* is the posterior probability of the null hypothesis is true given |*τ*| ≥ λ [16, 17, 18]:

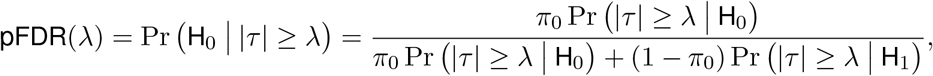

where H_0_ is the event that the null hypothesis is true, H_1_ is the event that the alternative hypothesis is true, and *π*_0_ is the proportion of true nulls. If there are *m* hypothesis tests then the expected number of false positives is *mπ*_0_ Pr (|*τ*| ≥ λ | H_0_) and the expected number of true positives is *m*(1 − π_0_) Pr (|*τ*| ≥ λ|H_1_). We can write the total number of expected significant genes as

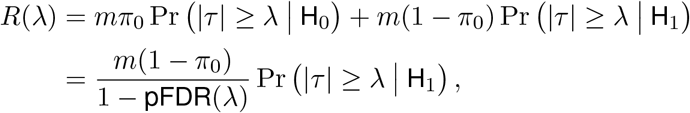

where we used the following relationship, *mπ*_0_ Pr (|*τ*| ≥ λ | H_0_) = *R*(λ)pFDR(λ). In order to maintain the pFDR(λ) at a fixed level when subsampling, the significance threshold varies in response to the changing power of the test statistics *τ* (*p*). Furthermore, as the read depth approaches infinity, there is an asymptotic number of significant genes 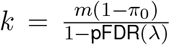. This value of *k* is not observed but it is the optimal number of discoveries when the technical variability is minimized with respect to the experimental design. The above equation can be rewritten in terms of the test statistic value and significance threshold at subsampling proportion *p* as

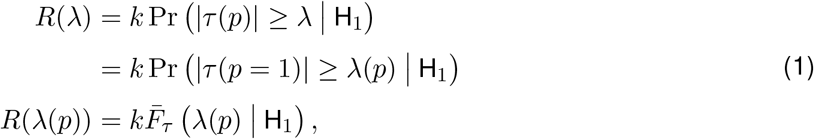

where *τ* (*p* = 1) are the observed test statistics from the completed experiment; 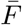 is one minus the cumulative distribution function (i.e., survivor function) of the test statistics under the alternative; and λ(*p*) is the changing significance threshold as a function of the subsampling proportion.

There are three unknowns in the above equation, namely, 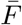, λ(*p*), and *k*. The cumulative distribution function of the test statistics under the alternative is unknown in practice. A related issue is the functional form of λ(*p*), which depends on the specific test statistic utilized. This is further complicated by any data transformations that were performed as part of the analysis. Finally, the asymptotic number of rejections *k* is unobserved. However, we can leverage the information from subsampling to approximate these unknown quantities using real data.

In order to model the relationship of the survival function, *R*(λ(*p*)), with respect to *p*, we take the derivative

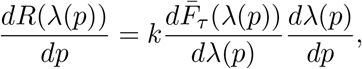

and empirically observe that the rate of change of *R*(λ(*p*)) has approximately a Log-normal distribution (Figure 4). As such, we assume that 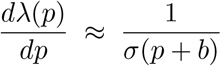 implying 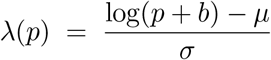, where (*μ*,*σ*) are the mean and standard deviation of the Normal distribution and *b* is an experiment-specific parameter. This gives our proposed model

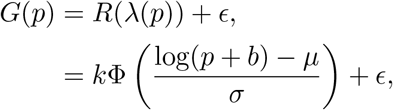

where *G*(*p*) are the observed number of significant genes, *R*(λ(*p*)) are the expected number of rejections, and *ϵ* are independent random errors from the model fit. The parameters *k*, *μ*, *b*, and *σ* in the probit model are estimated for each experiment using a non-linear least squares algorithm. It is important to note that these are not necessarily the parameters of the alternative distribution: this functional form is an approximation and the threshold parameter is unknown. We find that superSeq (solid line; Figure 4) provides a reasonable approximation to the observed empirical slopes.

As an alternative to modeling the survivor function using a Log-normal cumulative distribution function, we also considered a Logistic cumulative distribution function. In this case,

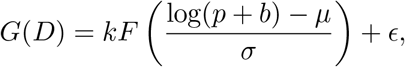

where 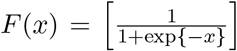. We fit this model, referred to as ‘logit’, using a non-linear least squares algorithm. Therefore, the logit and probit models are compared using the same number of free parameters: Figure 5 compares the parameter estimates under both models. As a baseline, we compared both of these models to a natural cubic smoothing spline with four degrees of freedom (referred to as the ‘smoother’ model).

**Figure 4:**
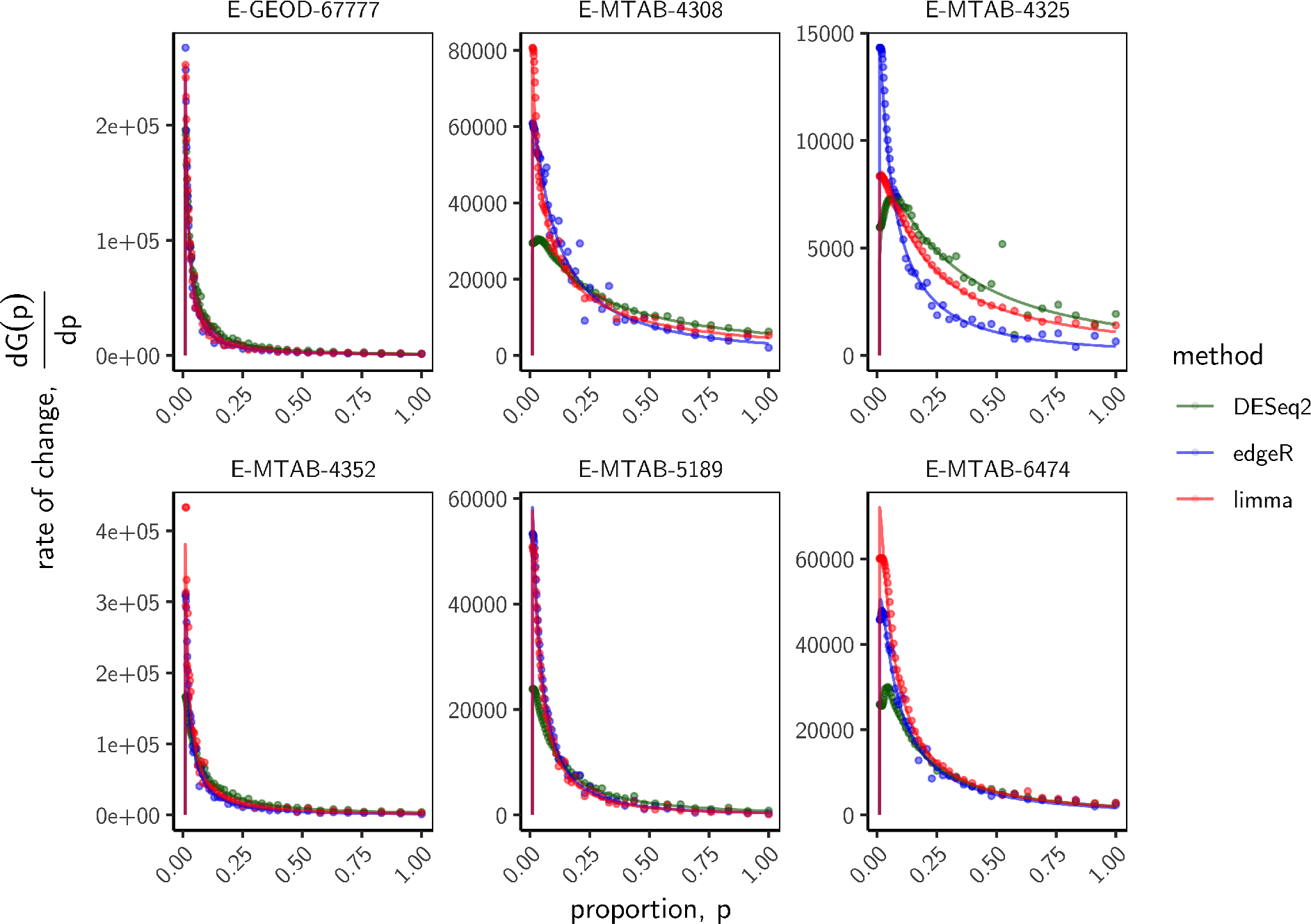
Observed slope in six randomly chosen experiments. For each experiment, we calculated the empirical slope at each subsampling proportion (denoted by the points) using the observed subsampling data. The solid lines represent the empirical slope from the predicted values of the superSeq model fit.

**Figure 5:**
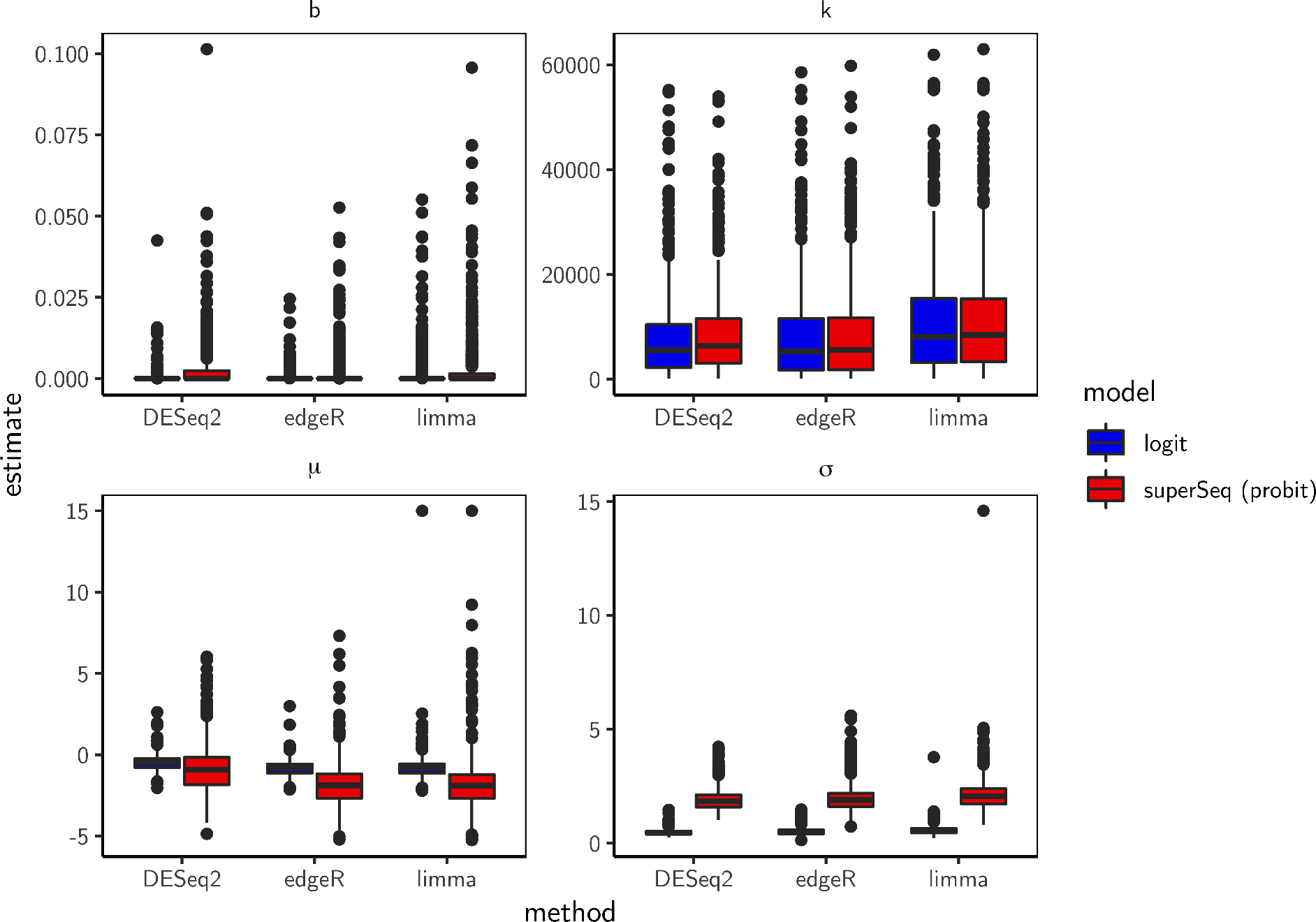
Distribution of parameter estimates in the superSeq model. The logit and probit model fits are shown using `DESeq2`, `edgeR`, and `limma`.

